# Intrusive Experiences In Post-Traumatic Stress Disorder: Treatment Response Induces Changes In The Effective Connectivity of The Anterior Insula

**DOI:** 10.1101/2020.10.01.319269

**Authors:** Arnaud Leroy, Etienne Very, Philippe Birmes, Sébastien Szaffarczyk, Renaud Lopes, Olivier Outteryck, Cécile Faure, Stéphane Duhem, Pierre Grandgenèvre, Frédérique Warembourg, Guillaume Vaiva, Renaud Jardri

**Affiliations:** Univ Lille, INSERM, CHU Lille, Lille Neuroscience & Cognition Centre (U-1772), Plasticity & SubjectivitY team, CURE platform, 59000 Lille, France; CHU Lille, Fontan Hospital, General Psychiatry Dpt., 59037 Lille cedex, France; Centre National de Ressources et Résilience pour les psychotraumatismes (CN2R Lille - Paris), 59000 Lille, France; CHU Toulouse, Purpan Hospital, Psychiatry Department, 31059 Toulouse Cedex, France; ToNIC, Toulouse NeuroImaging Center, INSERM U-1214, UPS, France; Univ Lille, INSERM, CHU Lille, Lille Neuroscience & Cognition Centre (U-1772), Degenerative & Vascular Cognitive Disorders team, 59000 Lille, France; Univ Lille, CNRS, INSERM, CHU Lille, Institut Pasteur de Lille, US 41 - UMS 2014 - PLBS, 59000 Lille, France; CHU Lille, Department of Neuroradiology, Roger Salengro Hospital, 59037 Lille cedex, France; CHU Lille, Fontan Hospital, Child & Adolescent Psychiatry Dpt., 59037 Lille cedex, France

**Keywords:** Post-Traumatic Stress Disorder, Treatment response, Effective connectivity, Salience network, fMRI, Reexperiencing

## Abstract

**Background:** One of the core features of posttraumatic stress disorder (PTSD) is reexperiencing the trauma. The anterior insula (AI) was proposed to play a crucial role in these intrusive experiences. However, the dynamic function of the AI in reexperiencing trauma, as well as its putative modulation by effective therapy, still need to be specified.

**Methods:** Thirty PTSD patients were enrolled and exposed to traumatic memory reactivation therapy. Resting-state fMRI scans were acquired before and after treatment. To explore AI directed influences over the rest of the brain, we referred to a mixed-model using pre/post Granger causality analysis seeded on the AI as a within-subject factor and treatment response as a between-subject factor. To further identify correlates of reexperiencing trauma, we investigated how intrusive severity affected: (i) causality maps and (ii) the spatial stability of other intrinsic brain networks.

**Results:** We observed dynamic changes in AI effective connectivity in PTSD patients. Many within- and between-network causal paths were found to be less influenced by the AI after effective therapy. Insular influences were found positively correlated with flashback severity, while reexperiencing was linked with a stronger *default mode network* (DMN) and more unstable *central executive network* (CEN) connectivity.

**Conclusion:** We showed that directed changes in AI signaling to the DMN and CEN at rest may underlie the degree of intrusive symptoms in PTSD. A positive response to treatment further induced changes in network-to-network anticorrelated patterns. Such findings may guide targeted neuromodulation strategies in PTSD patients not suitably improved by conventional treatment.

## INTRODUCTION

Posttraumatic stress disorder (PTSD) is a disabling condition that can be triggered by terrifying events that have the potential to disrupt life, such as interpersonal violence, combat, life-threatening accidents or disasters, as well as global pandemics (Horesh and Brown, 2020). PTSD may lead to chronic psychiatric or addictive morbidities, loss of normal daily functioning, and increased risk of suicide (Lewis et al., 2019). This disorder usually induces intrusive symptoms (i.e., distressing recollections of the event, including flashbacks and nightmares, often called ‘‘re-experiences’’), persistent avoidance of stimuli associated with the trauma, negative alterations in cognitions or mood, and hyperarousal (American Psychiatric Association, 1994). ‘‘Re-experiencing’’ is considered central in the pathophysiology of PTSD, despite some similarities with other intrusive thoughts observed transdiagnostically, such as hallucinations, ruminations or persistent worries (Brewin et al., 2010; Larøi et al., 2012; Newman et al., 2013; Watkins, 2008). Even if this research field is prolific, it still lacks a common neurofunctional signature for intrusive experiences that adequately circumscribes the underlying mechanisms of PTSD.

Brain-wide dysfunctions have already been suggested at the root of PTSD. First fMRI evidence came from task-based studies, in which structures involved in memory and emotional processing (e.g. amygdala, hippocampus or ventral prefrontal cortex), were reported as key elements of the neurocircuitry of PTSD (Mahan and Ressler, 2012). Interestingly, another candidate node – the anterior insula (AI) - picked the interest of trauma-focused scientists beyond fear-processing. The bilateral AI was indeed repeatedly found overactive in PTSD, from women exposed to intimate-partner violence (Fonzo et al., 2010), to veterans (Duval et al., 2020). Crucially, activation-level in the insula and its connected structures, such as the amygdala (Yehuda et al., 2015) and the hippocampus (Stevens et al., 2018), was found associated with hyperarousal and re-experiencing, suggesting a specific role of the AI in these clinical dimensions.

On a more general level, the AI appears as one of the major connector hubs in the brain (van den Heuvel and Sporns, 2013). This structure has been implicated in a large variety of physiological functions, ranging from feelings representation to bodily and self-awareness (Gogolla, 2017). It receives convergent inputs from multiple sensory modalities, including the auditory and visual systems (Augustine, 1996; Bamiou et al., 2003; Butti and Hof, 2010; Mesulam and Mufson, 1982; Nieuwenhuys, 2012), while converging evidence supports AI’s involvement in simultaneous attention to multisensory events (Bushara et al., 2003, 2001). The AI was also proposed to tag salient endogenous and external information and further reallocate attentional resources towards them (Menon and Uddin, 2010), making it a central element of the ‘‘salience network’’ (SN).

Besides task-based fMRI studies, a second line of evidence about the role of AI came from correlational mapping between a seed and other regions of the brain, also called functional connectivity or FC approaches. These studies allowed to explore how these areas cross-talk and potentially how they relate to clinical dimensions of PTSD. Again the AI was found particularly involved. An increased FC was notably reported at rest between the bilateral insula with the lingual gyri and precuneus (both involved in implicit memory processing) in the dissociative sub-type of PTSD (Harricharan et al., 2020). These resting-state dysconnectivity patterns were found to be state-dependent, and susceptible to change when subjects with PTSD receive adequate treatments.

For instance, following psychotherapy, the AI and amygdala exhibit increased reciproqual connectivity, but also an increased FC with the ventral prefrontal cortex, frontopolar and sensory cortices, while other regions, such as the left frontoparietal nodes of the central executive network, rather show decreased FC at rest (Fonzo et al., 2021). This is also the case for larger decreases in amygdalar–frontal connectivity and AI–parietal connectivity, both found associated with PTSD symptom reductions (Fonzo et al., 2021). These results suggest that subtle interactions between AI and the brain regions involved in cognitive control or emotional processing are associated with treatment response in PTSD.

Despite indisputable progress, we can notice that only a limited number of studies explored more in detail the fine-grained influence that AI exerts over other structures. A first study, conducted within the SN, evidenced a reduced dynamical causal flow from the right amygdala to the right insula (Weng et al., 2019), while pivotal changes in connectivity strength and temporal variability between the right AI and the middle frontal gyri were also measured (Rangaprakash et al., 2018a), again supporting a key role for AI in PTSD. However, these studies neither assessed intrusive symptoms (and only one explored treatment response (Fonzo et al., 2021)), nor the whole-brain effective connectivity of the AI, since they limited the analysis to a pre-defined set of regions of interest. This approach may appear in contradiction with the well-accepted view that the brain is an interconnected network of functional components rather than made of discrete units. The AI, and by extension the SN, appears to be one of these functional components.

We can illustrate this view looking at intrusive symptoms more broadly. SN effective connectivity was indeed proven involved in the process of switching from a state of unconstrained rest to one of experiencing hallucinatory events (Lefebvre et al., 2016; Palaniyappan and Liddle, 2012), reinforcing the hypothesis that this network may govern intrusive experiences in general. The SN is thus thought to tightly control the balance between various intrinsic networks and to swiftly move from rest to task-based actions and vice-versa, a theory that has been conceptualized as the tripartite model (Koch et al., 2016; Lefebvre et al., 2016; Menon and Uddin, 2010; Sridharan et al., 2008; Stevens et al., 2018; Yehuda et al., 2015). According to this framework, the SN may drive commonly observed anticorrelated patterns between the default mode network (DMN, underlying self-referential processing) and the central executive network (CEN, involved in cognitive control and decision making) (Menon and Uddin, 2010), an interaction already shown to be impaired in PTSD. Interestingly, surges in connectivity strength of the CEN were reported to be associated with intrusiveness (Koch et al., 2016; Nicholson et al., 2016; Sheynin et al., 2020; Weng et al., 2019).

Even encouraging, some gaps remain in our understanding of the exact network dynamics underlying intrusive experiences in PTSD and their resolution. We notably still lack causal proofs about the directionality of the neural alterations underlying intrusive symptoms in PTSD. The present study intends to bring new insights to the dynamic role of the SN in PTSD and, more specifically, to circumscribe the directed influence of AI over the rest of the brain as a function of treatment response. To do so, we analyzed fMRI data from a recent randomized controlled trial (RCT) comparing trauma memory reactivation therapy using propranolol (considered a putative reconsolidation blocker) versus the same therapy plus placebo in PTSD patients (Roullet et al., 2021). Similar designs previously showed a significant symptom severity reduction after joint therapy (Brunet et al., 2018). Surprisingly, the most recent RCT did not confirmed superiority for joint therapy with propranolol over therapy plus placebo (Roullet et al., 2021), but a significant proportion of patients was nevertheless improved in both groups. We took this opportunity to explore dynamical brain markers of treatment response and decided to focus on re-experiencing symptoms, as one possible cause of limited treatment efficacy may rely not only on treatments themselves, but also in the heterogeneity within PTSD or selective actions on clinical dimensions and not others (Neria, 2021).

Here, we hypothesize that effective therapy could modulate the effective connectivity of AI and that these plastic changes correlate with a reduction in trauma re-experiencing symptoms. In reference to the tripartite model, we also expect this downgrading in intrusiveness to be linked to changes in DMN and CEN spatial stability, demonstrating a brain-wide reallocation of cognitive resources in PTSD patients who respond to treatment.

## MATERIALS AND METHODS

### Population

Patients with a primary diagnosis of PTSD according to DSM-IV-TR criteria (Structured Clinical Interview for DSM-IV, PTSD module), and in the absence of contraindication to propranolol, provided written consent to participate in the research (Main characteristics and inclusion/exclusion criteria are reported in **Table 1**). They all took part in a RCT testing for the efficacy of traumatic memory reactivation under the influence of propranolol versus placebo. This trial received approval from an ethics committee (CPP 2009-012976-29) and was registered on clinicaltrials.gov (NCT01713556). The main results for this clinical trial (i.e., that the efficacy of propranolol was not greater than that of placebo one week posttreatment) are presented elsewhere (Roullet et al., 2021).

**Table 1:** Population description.

|  | Responders (n=16) | Nonresponders (n = 14) |
| --- | --- | --- |
| Age | 41.3 (12.3) | 36.6 (13.5) |
| Sex (Male /Female) | 7/9 | 8/6 |
| Dvars | 24.3 (3.74) | 24.1 (4.05) |
| Global signal | 887 (121) | 916 (96.3) |
| Total PCL-S score | 65.4 (9.34) | 68.4 (9.34) |
| PCLS-score (question 1 to 5) | 18.9 (3.38) | 19.4 (3.38) |
| Beck Depression inventory score | 25.9 (14.5) | 28.9 (11.3) |
| *Rate of patients receiving propranolol \** | 31.3 % | 78.6% |

\*p <0.05. Inclusion and exclusion criteria: primary diagnosis of PTSD according to DSM-IV-TR criteria (Structured Clinical Interview for DSM-IV, PTSD module), absence of contraindication to propranolol (hypotension, higher than a first-degree heart block, bronchial asthma etc.); medication recommended or suggested for PTSD treatment; psychotherapy; basal systolic blood pressure < 100mm Hg; basal heart rate <50 bpm; psychotic or bipolar disorders; traumatic brain injury; current substance or alcohol dependence; acute suicidal ideation; pregnancy and breast feeding.

Participants waiting for treatment were randomly and blindly allocated to two experimental groups (1:1 ratio): (i) a ‘‘traumatic memory reactivation therapy + propranolol’’ group, and (ii) a ‘‘traumatic memory reactivation therapy + placebo’’ group. Propranolol or placebo was administered 90-minutes before the memory reactivation session, performed once a week for 6 consecutive weeks. The *Posttraumatic Stress Disorder Checklist Scale* - PCL-S (Ventureyra et al., 2002) was used to quantify symptom severity and assess treatment response. Because we were interested in intrusive symptoms, we focused on the item sum of PCL-S Q1-to-Q5 (Ventureyra et al., 2002). Assessment was made before treatment (at baseline) and one week after the end of the treatment (post-treatment). Patients had brief memory reactivation session with or without propranolol, once a week for 6 consecutive weeks. The response to memory reactivation (with or without propranolol) in the whole sample (sometimes referred to as therapy in this paper for the sake of simplicity (Thierrée et al., 2020)) was considered positive for at least a 33% decrease (Brady et al., 2015; Mushtaq et al., 2012) in the PCL-S Q1-to-Q5 score compared to baseline. This threshold is commonly referred to as the first significant level of response also called ‘‘poor response’’.

### MRI acquisition and data preprocessing

Patients underwent two MRI sessions at rest with their eyes closed (at baseline and post-treatment) on a 3T Philips Achieva scanner with an 8-channel head coil. Each of these sessions included a 4-min T1-weighted (T1w) 3D anatomical run (124 transverse slices, field of view = 256 mm^3^, vox = 0.8 mm^3^) and a 15-min T2*-weighted 3D-PRESTO sequence (Liu et al., 1993; Neggers et al., 2008; van Gelderen et al., 2012). This functional sequence (dynamic scan time = 1000 ms, TE = 9.6 ms, flip angle = 9°, vox = 3.3 mm^3^) allowed for full functional brain coverage with a temporal resolution particularly suited for effective connectivity analysis (Deshpande and Hu, 2012).

Anatomical and functional MRI data were preprocessed using the FMRIPrep pipeline v. 1.5 (Esteban et al., 2019), a Nipype v. 1.2.2 based tool (Gorgolewski et al., 2011). T1w images were corrected for nonuniform intensity and skull stripped. Brain tissue segmentation of cerebrospinal fluid, white-matter and gray-matter was performed on the brain-extracted T1w image. Volume-based spatial normalization to the *Montreal Neurological Institute* ICBM-152 (MNI) template was performed through nonlinear registration using brain-extracted versions of both the T1w reference and the MNI template.

For functional images, a reference BOLD volume and its skull-stripped version were generated and coregistered to the T1w image using a boundary-based registration algorithm with 9 degrees of freedom. Head-motion parameters were estimated before spatiotemporal filtering. Motion correction, BOLD-to-T1w transformation and T1w-to-template (MNI) warps were concatenated and applied in a single step using antsApplyTransforms (ANTs v2.1.0) based on Lanczos interpolation. Spatial smoothing with a 6-mm isotropic Gaussian kernel was then performed, and a second “nonaggressive” denoising step was conducted using independent component analysis [ICA-AROMA]. Linear trends were finally removed, and a high-pass temporal filtering with 3 cycles/point was applied.

### Granger causality analysis

Effective connectivity was assessed using Granger causality analysis (GCA), which allows for the data-driven exploration of a reference region’s influence over the brain as well as targets of influence on the same given area (Roebroeck et al., 2005). We used this approach to account for the multiple regions naturally connected to the SN (Menon, 2015) and to avoid missing brain areas that could not have been retained in a more traditional theory-driven framework. Here, we used the implementation proposed in the BrainVoyager software suite (v21.4, BrainInnovation, Maastricht; *Granger Causality Mapping* plugin V1.5). We defined the bilateral AI, according to the meta-analysis from Laird et al. (**Figure 1**) (2011) (Laird et al., 2011), as the reference region for later analyses. GCA maps were used to visualize the directed influences between the AI and every voxel in both directions after applying a gray-matter mask. These maps were thresholded using a false-discovery rate approach at q-levels of 0.01. We finally tested the association between GCA maps and symptom severity at the “post-treatment” time-point, considering p_fwe_<0.05 as significant.

**Figure 1.**
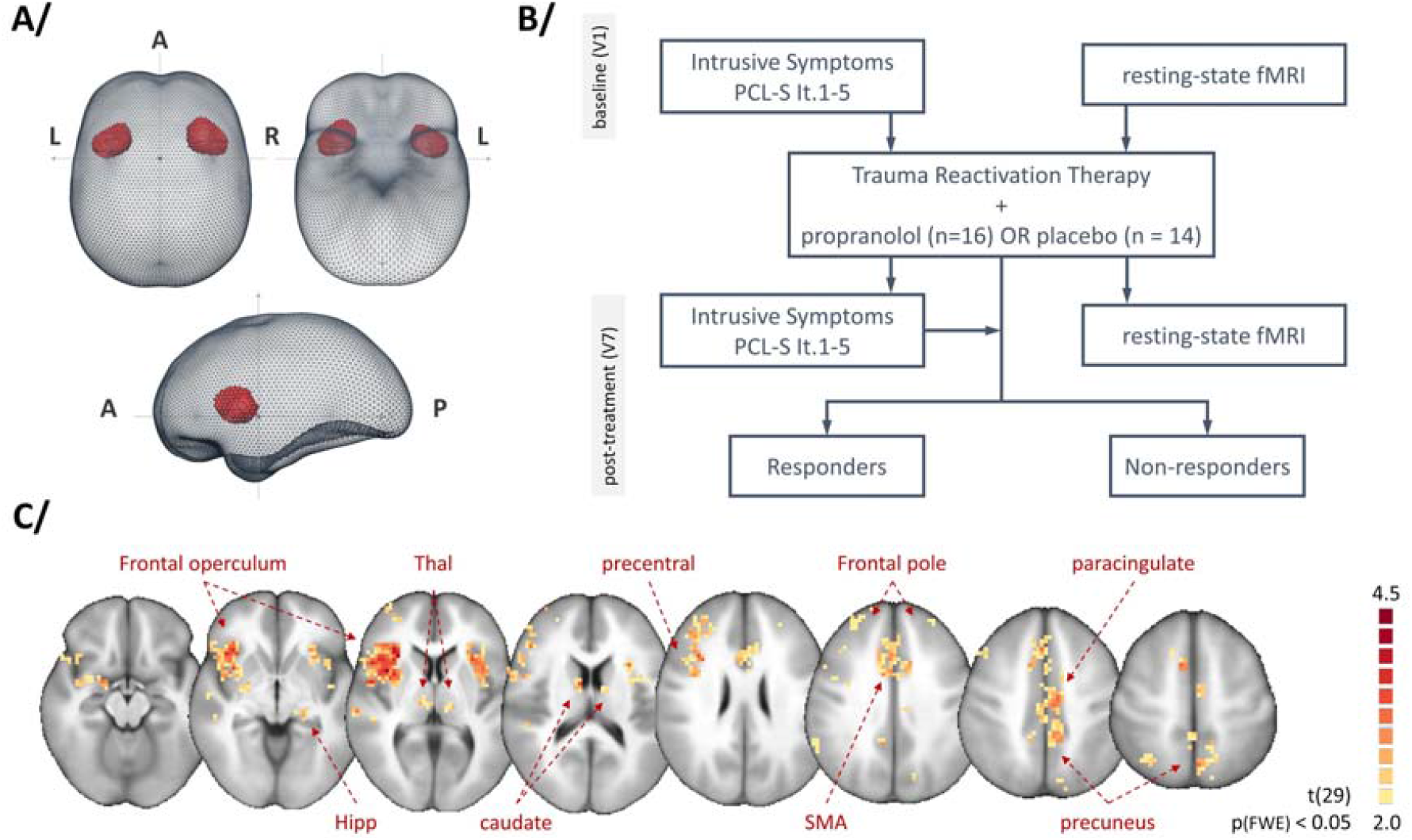
Study design and Granger causality analysis (GCA) seeded on the anterior insula in PTSD patients. **(A)** The bilateral anterior insula was chosen as the region of interest for GCA and is presented in red in a glass brain. **(B)** Flow chart of the study. We defined responders as patients with at least a 33% decrease in PCL-S scores post-treatment compared with baseline. **(C)** Whole-sample random-effects GCA map at baseline. PCL-S: Posttraumatic Stress Disorder Checklist Scale; Thal: thalamus; Hipp: hippocampus; SMA: supplementary motor area; A: anterior; P: posterior; L/R: left/right sides of the brain.

### Main statistical analysis

To assess changes in AI effective-connectivity pre/post treatment between patients who respond to treatment (responders) and those who did not (nonresponders), we referred to a 2×2 mixed-model ANOVA, using GCA maps at baseline and post-treatment as within-subject factors and treatment response as a between-subject factor. Post-hoc analyses used Student’s t-tests. All maps were then thresholded using a cluster-based permutation method (Forman et al., 1995). To prevent potential inflated false-positive rates (Eklund et al., 2016), we first specified a cluster-defining threshold (CDT) at p_uncorrected_ < 0.001. After conducting a 1000-iteration Monte-Carlo simulation, a cluster-extent threshold was defined as a value high enough to keep the family-wise error at p_fwe_=0.05. The resulting brain areas were labeled using the Anatomy toolbox v 3.0 (https://github.com/inm7/jubrain-anatomy-toolbox (Eickhoff et al., 2005)).

### Intrinsic network spatial stability measure

In parallel, we also explored the spatial stability of the DMN and the CEN posttreatment (i.e., the “post-treatment” time-point) using a ‘‘goodness-of-fit’’ (GoF) procedure. After decomposing each posttreatment functional dataset using *independent component analysis* (ICA), we selected the components exhibiting the highest spatial correlation with an a priori template. For each participant, this procedure was repeated twice: with the DMN and the CEN template, respectively (Greicius et al., 2004). The resulting GoF scores were assumed to reflect posttreatment DMN and CEN spatial stability. We tested for an association between these GoF scores and the severity of intrusive symptoms using Pearson’s *r* correlation test, considering p<0.05 as significant.

## RESULTS

### Demographic and clinical variables

Among the initial sample of 66 participants, 59 completed the treatment and 30 performed the full pre/post fMRI assessment. At baseline, responders and nonresponders were comparable in terms of (i) sociodemographic characteristics (age and sex-ratio), and (ii) symptom severity (PCL-S scores/subscores and depressive symptoms measured with the *Beck Depression Inventory* (Beck et al., 1988)). Note that the responder group contained fewer patients receiving propranolol (*Table 1*, see (Roullet et al., accepted) for a more detailed description of the RCT main findings).

### AI effective connectivity before treatment

GCA performed on the whole-sample at baseline revealed that the bilateral AI significantly modulates a group of regions involved in motor preparation, execution and action monitoring (i.e., precentral gyrus, the supplementary motor area, the left thalamus, and the frontal pole), as well as in visuospatial processing (i.e., the paracingulate gyrus and the precuneus). See **Figure 1** and **Suppl. Table 2**. No specific connectivity differences were evidenced between the responder and nonresponder groups before treatment.

### Pre/post AI effective-connectivity changes

The mixed-model ANOVA revealed significant changes in AI causal maps after treatment. A significant [time-point x group] interaction was evidenced: (i) laterally, in the mid- and posterior insula, amygdala, precentral gyrus and supramarginal gyrus; and (ii) medially, in the cingulate cortex (anterior and posterior) and the precuneus. Compared with nonresponders, patients showing clinical improvement exhibited a reduced influence of the AI over a wide socioemotional network composed of the superior frontal gyrus, anterior and posterior supramarginal gyri, anterior and posterior cingulate, central operculum and right amygdala. Conversely, responders exhibited a higher influence of the AI over the precuneus (Cf. **Figure 2, Table 2**). As a control, we repeated this analysis using the left (or right) anterior insula as seed, and evidenced a high-degree of overlapping regions between the bilateral insula and the right insula seeds (left supramarginal gyrus, right central opercular cortex, right putamen), and between the bilateral insula and the left insula seeds (cingulate gyrus, anterior division, right precuneus) (**See Suppl. Figure 1**). Simple pre/posttreatment contrasts for responders and nonresponders are available in **Suppl. Figures 2, 3** and **Suppl. Tables 3, 4** respectively, As a point of comparison, significant results for propranolol vs placebo after treatment are presented in **Suppl. Figure 4 and Suppl. Table 5**.

**Figure 2.**
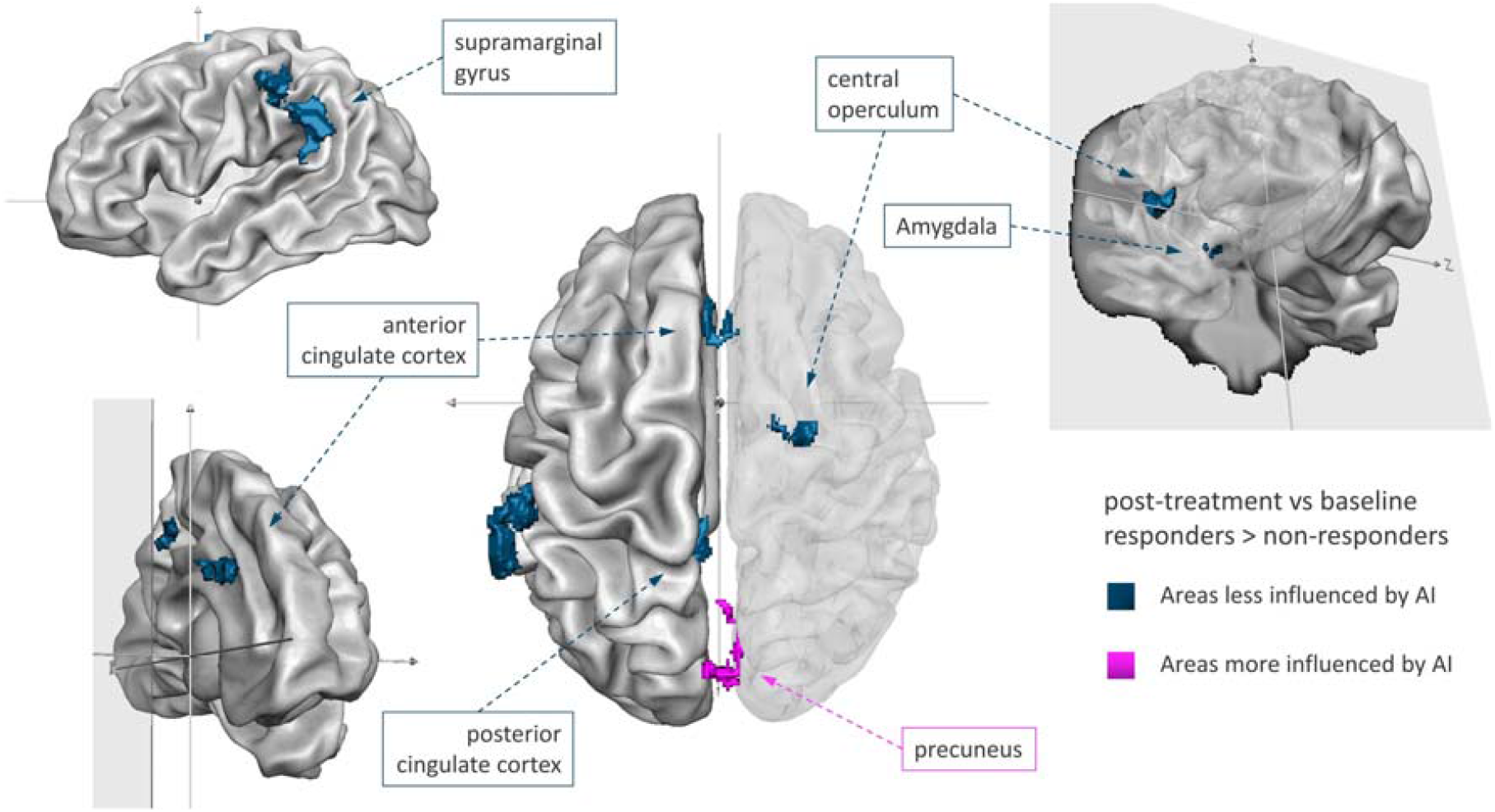
Changes in Granger causality maps seeded on the anterior insula (AI) between responders and non-responders to therapy in PTSD. We used a transparent right hemisphere to allow visualization of the deeper clusters. Brain areas less influenced by AI after effective treatment are depicted in dark blue. The precuneus (pink) was the only cluster found to be more influenced by AI posttreatment in responders than in non-responders.

**Table 2:** Changes in the anterior insula effective connectivity between responder and non-responder patients. Regions exhibiting a significant difference in *Granger causality analysis* maps between baseline and post-treatment according to treatment response. Coordinates are reported in the MNI (Montreal Neurological Institute) space.

| x | y | z | t-value | p-value | Label |
| --- | --- | --- | --- | --- | --- |
| -63 | -45 | 36 | -3.813837 | 0.000662 | Left Supramarginal Gyrus.<br>posterior division |
| -57 | -30 | 39 | -4.072859 | 0.000328 | Left Supramarginal Gyrus. anterior<br>division |
| -6 | -39 | 45 | -3.452251 | 0.001728 | Cingulate Gyrus. posterior division |
| -3 | 30 | 27 | -2.993126 | 0.005594 | Cingulate Gyrus. anterior division |
| 6 | -72 | 42 | 3.098338 | 0.004296 | Right Precuneus Cortex |
| 21 | -9 | 66 | -3.458369 | 0.0017 | Superior Frontal Gyrus |
| 27 | 3 | -12 | -2.894758 | 0.007138 | Right amygdala |
| 45 | 0 | 9 | -2.875736 | 0.007479 | Right Central Opercular Cortex |

Regression analysis conducted posttreatment further revealed that the more severe the intrusive symptoms were, the greater the AI exerted influence over somatosensory and motor regions (i.e., the posterior insula, right parietal operculum, and precentral gyrus), as well as over brain areas involved in visuospatial processing (the paracingulate gyrus) and self-other processing (i.e., the anterior and posterior cingulate cortices, superior frontal gyrus and supramarginal gyrus). See **Figure 3**.

**Figure 3.**
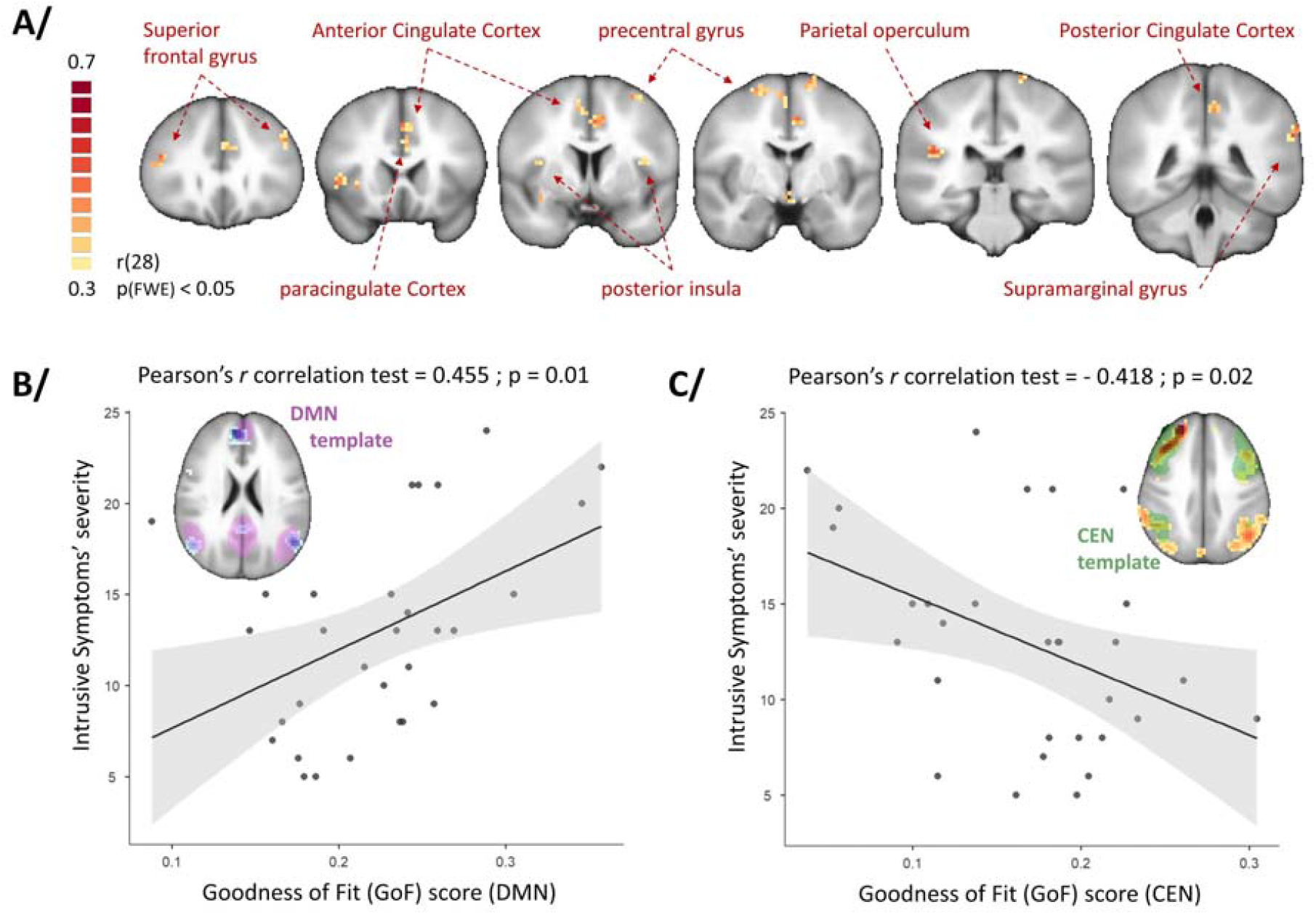
Brain correlates of intrusive symptom severity in PTSD. **(A)** Linear regression analysis showing the brain regions exhibiting a positive association between the severity of intrusive symptoms and thresholded *Granger causality analysis* maps post-treatment. **(B, C)** Correlation analyses between intrinsic network stability and the severity of intrusive symptoms post-treatment. A positive association was indicated by the default mode network stability (DMN GoF score), shown in **(B)**, whereas a negative association was indicated by the central executive network stability (CEN GoF score), shown in **(C)**.

### Intrinsic network spatial stability measure after treatment

Finally, the assessment of the spatial stability of the DMN and CEN posttreatment revealed that the severity of ‘‘re-experiencing’’ symptoms was positively correlated with the DMN GoF scores (**r = 0.521, p = 0.003**) and negatively correlated with the CEN GoF scores (**r = −0.418 p = 0.021** - Cf. **Figure 3**).

## DISCUSSION

The present fMRI study was designed to explore how an effective response to traumatic memory reactivation therapy (with or without propranolol) for intrusive symptoms in PTSD could modulate AI causal influences over the brain. Many fMRI studies measured pre/post functional connectivity using correlations (Etkin et al., 2019; Korgaonkar et al., 2020; Santarnecchi et al., 2019; Zhu et al., 2018), but only a limited number of studies reported effective connectivity changes in PTSD (Fonzo et al., 2021; Rangaprakash et al., 2018b), with no particular focus on intrusive symptoms in a whole brain approach. We focused on the AI since this region is known to be: (i) a central integration hub serving sensory, emotional, motivational and cognitive functions, (ii) potentially involved in re-experiencing trauma. Even if the native RCT was not specifically design to test for the prediction of response, we were able, using high temporal-resolution fMRI and comparing pre/post therapy GCA results in PTSD patients, to provide new evidence that treatment response was associated with a significant reduction in AI effective connectivity towards motor and socioaffective regions, a global decrease that follows symptom severity reduction.

The first set of regions modulated by AI corresponded to limbic areas for which dense reciprocal connections with the ventral AI were repeatedly described. A strong body of evidence supports AI mediation in fear and anxiety, which was regularly found to be coactivated with the amygdala in stressful contexts (Gogolla, 2017). By showing a reduced influence of AI on the amygdala in responders compared with nonresponders, we can assume that one of the first effects of treatment was to temper the emotional storm associated with re-experiencing and hyperarousal (Yehuda et al., 2015). This finding nicely complements the existing literature in which joint amygdalar and insular overactivation is described in the context of PTSD (Koch et al., 2016) and declines after successful therapy (Malejko et al., 2017). The ability to better modulate AI connectivity following therapy could be associated with better cross-talk between untargeted inner thoughts and the ability to focus attention on stimulus-dependent demands (Szeszko and Yehuda, 2019), a theory also supported by the association found between symptom severity and AI influence over visuospatial areas. Using GCA, we were able to attest that AI primarily drives this pathological interaction in PTSD.

The second set of brain areas modulated by AI are involved in self-other distinction and may support dissociative experiences frequently observed in PTSD. This is the case for the precuneus, frontal superior and supramarginal gyri, all regularly found to be involved in self-awareness and agency processing (e.g., (Sperduti et al., 2011)). These cognitive functions are more usually under the influence of the dorsal AI (Kurth et al., 2010). Interestingly, localized AI lesions can induce dissociative experiences, such as the (rare) “pain asymbolia” syndrome, in which pain recognition appears disconnected from its appropriate emotional response (9). Within this functional network, the supramarginal gyrus, located at the temporo-parietal junction, has also been linked with experiences involving a sensorial component. Similar to the AI, the supramarginal gyrus receives heavy sensory inputs ranging from the auditory to the somatosensory modality. The crossmodal nature of this area makes it particularly well suited for linking sensory experiences with cognitive and/or affective information. Finally, the supramarginal gyrus is also involved in the phonological and articulatory processing of words (Stoeckel et al., 2009), making it solicitable by talking therapy. Again, this is perfectly in line with the present findings showing that the AI influence on this network was correlated with the degree of intrusion and was significantly decreased in responders.

The third set of regions influenced by AI is engaged in sensorimotor control (Chouinard and Paus, 2006; Dum and Strick, 1991; Paus, 2001) and might be involved in the autonomic and behavioral responses to stress. Again, this interaction was found to correlate with symptom severity, even if only indirectly through the posterior insula and thalamic relays (Uddin et al., 2017), which were found to be under AI control at baseline. The motor network under consideration includes the caudal anterior cingulate cortex (cACC), the supplementary motor area and the precentral gyrus. A decreased resting-state functional connectivity between the cACC and the precentral gyrus was previously evidenced in veterans with or without PTSD compared to healthy controls, suggesting that military training or deployment, including trauma exposure, may influence SN connectivity (Kennis et al., 2014). In addition, precentral activity has been related to defensive behaviors in animals (Graziano and Cooke, 2006) and may subserve the increased “fight or flight” response regularly observed in PTSD when facing mental stress.

In addition to networks sustaining the rich phenomenology of PTSD symptoms, we also investigated how effective treatment may dynamically affect the interaction between intrinsic neural networks. We first demonstrated that the more the AI exerted a causal influence over core nodes of the DMN (such as the rostral ACC, the posterior cingulate and the precuneus), the more severe intrusive symptoms were, and that this interaction differentially changed according to the treatment response. This finding appears in line with previous studies conducted in schizophrenia patients (Lefebvre et al., 2016), showing that increased control from the SN to the DMN initiates hallucinatory states (another example of intrusive experiences). Interestingly, the DMN is also known to anti-correlate with task-related networks, such as the CEN (Fox et al., 2005; Greicius et al., 2003), and this antagonistic activity was proposed to be tuned by the AI (Sridharan et al., 2008). Returning to our schizophrenia example, a CEN takeover was found to drive the extinction of hallucinations (Lefebvre et al., 2016). Here, we found that re-experiencing the trauma positively correlated with DMN stability and presented a reversed pattern for the CEN. Altogether, these results support the idea that intrusive symptoms could correspond to self-referential mental activities driven by impaired AI control over the DMN/CEN balance, making salient memory fragments active enough (through bottom-up amplification) to aberrantly intrude into consciousness.

Despite these encouraging findings, some issues need to be further discussed. If the above-mentioned theory is correct, we could expect to find the hippocampal complex among the regions influenced by the AI. The limited sample-size of the present trial may account for such a negative result (i.e., a power issue), and the exact relationship between the AI and this limbic structure will have to be clarified in future studies. In fact, several brain areas identified in this study have previously been shown to be involved in memory suppression beyond the medial temporal lobe. This is the case for the precuneus and the frontal cortex (Mary et al., 2020), which have tight connections with the hippocampus (Anderson et al., 2016; Cunningham et al., 2017). The same is true for other limbic structures, such as the amygdala which have strong reciprocal connections with the hippocampus. We cannot exclude that intermediate small brain structures, such as the hippocampus, that are involved in a chain of causality could be more vulnerable and may not survive statistical thresholding. Based on that possibility, we hypothesize a triple interaction [AI - amygdala - hippocampus] at the root of the memorization of trauma-related emotional valence, constituting an interesting complementary track for future research on PTSD.

A second potential issue resides in the fact that GCA indicates the dominant direction of influence, introducing ambiguity in the interpretation of pre/post treatment contrast maps, as they may potentially result from a decrease in the influence of AI-to-target-regions influence or from an increase in the influence of target-to-AI. This problem can, for instance, be illustrated by considering a recent study of effective connectivity in PTSD that reached seemingly opposite conclusions, suggesting that frontal regions exerted a reduced influence on AI (Rangaprakash et al., 2018a). In the same vein, a decreased causal flow from the right amygdala to the right insula in PTSD patients relative to trauma-exposed controls has been suggested (Weng et al., 2019), which again appears in apparent contradiction with the present findings. Even if the samples and designs were not exactly comparable, methodological advances should help to reconcile these findings, but until then, this literature needs to be interpreted with caution and in reference to clinically and anatomo-functionally available knowledge at the time of publication.

A final point we would like to insist on is that we focused on symptom reduction, regardless of the initial group of randomization in the *Pre-Reactivation Propranolol Therapy* trial. Of course, the full results of the RCT have been presented elsewhere (Roullet et al., accepted) and are beyond the scope of the present paper. Note that the between-group differences in our study were both explained by changes in the responder and non-responder groups. This is compatible with previous studies showing that psychotherapy could induce functional changes, even in cases of non-significant clinical response (Simmons et al., 2013), that could be linked with repeated trauma exposure without extinction, as suggested in previous studies (Uddin et al., 2017). Importantly, because half of the sample did not reach the threshold for a positive response to therapy, we expect our findings to also be relevant for future neuromodulation trials for severely impaired PTSD patients. Neuromodulation methods such as transcranial magnetic stimulation usually targets superficial cortical areas, and AI could constitute a better target for neurofeedback which allows to modulate the activity/connectivity of profound or sub-cortical structure. Promisingly, a recent study confirmed that PTSD patients could be trained to downregulate amygdalar activity using real time fMRI-based neurofeedback (de Pierrefeu et al., 2018).

Overall, we were able to provide experimental support for the causal role played by AI in re-experiencing trauma. Notably, we showed that effective therapy was linked with plastic changes in AI directed influence over sensorimotor, cognitive and socioemotional networks. Dynamically, restoration of the DMN-to-CEN switch control was also observed, offering an attractive mechanism for intrusion. We hope that the present results will contribute to paving the way for new evidenced-based treatments of intrusive symptoms in PTSD, considering AI as a particularly interesting target for this purpose.

## ACKNOWLEDGMENTS

Funding source: French Ministry of Health’s Hospital Program of Clinical Research.

## AUTHOR CONTRIBUTIONS

AL, EV, PB, SS, RL, CF, SD, PG, FW, GV, RJ contributed to the conception and design of the work AL, SS, RL, RJ contributed to the interpretation of data for the work

AL, EV, PB, GV, RJ drafted the work

AL, EV, PB, SS, RL, CF, SD, PG, FW, GV, RJ gave the final approval of the version to be published

## REFERENCES

American Psychiatric Association, 1994. Diagnostic and Statistical Manual of Mental Disorders, Fourth Edition. ed. American Psychiatric Association.

Anderson, M.C., Bunce, J.G., Barbas, H., 2016. Prefrontal-hippocampal pathways underlying inhibitory control over memory. Neurobiol Learn Mem 134 Pt A, 145–161.

Augustine, J.R., 1996. Circuitry and functional aspects of the insular lobe in primates including humans. Brain Research Reviews 22, 229–244.

Bamiou, D.-E., Musiek, F.E., Luxon, L.M., 2003. The insula (Island of Reil) and its role in auditory processing: Literature review. Brain Research Reviews 42, 143–154.

Beck, A.T., Steer, R.A., Carbin, M.G., 1988. Psychometric properties of the Beck Depression Inventory: Twenty-five years of evaluation. Clinical Psychology Review 8, 77–100.

Brady, F., Warnock-Parkes, E., Barker, C., Ehlers, A., 2015. Early in-session predictors of response to trauma-focused cognitive therapy for posttraumatic stress disorder. Behav Res Ther 75, 40–47.

Brewin, C.R., Gregory, J.D., Lipton, M., Burgess, N., 2010. Intrusive images in psychological disorders: characteristics, neural mechanisms, and treatment implications. Psychol Rev 117, 210–232.

Brunet, A., Saumier, D., Liu, A., Streiner, D.L., Tremblay, J., Pitman, R.K., 2018. Reduction of PTSD Symptoms With Pre-Reactivation Propranolol Therapy: A Randomized Controlled Trial. Am J Psychiatry 175, 427–433.

Bushara Grafman, J., Hallett, M., 2001. Neural correlates of auditory-visual stimulus onset asynchrony detection. J. Neurosci. 21, 300–304.

Bushara Hanakawa, T., Immisch, I., Toma, K., Kansaku, K., Hallett, M., 2003. Neural correlates of cross-modal binding. Nat. Neurosci. 6, 190–195.

Butti, C., Hof, P.R., 2010. The insular cortex: a comparative perspective. Brain Struct Funct 214, 477–493.

Chouinard, P.A., Paus, T., 2006. The primary motor and premotor areas of the human cerebral cortex. Neuroscientist 12, 143–152.

Cunningham, S.I., Tomasi, D., Volkow, N.D., 2017. Structural and functional connectivity of the precuneus and thalamus to the default mode network. Hum Brain Mapp 38, 938–956.

de Pierrefeu, A., Fovet, T., Hadj-Selem, F., Löfstedt, T., Ciuciu, P., Lefebvre, S., Thomas, P., Lopes, R., Jardri, R., Duchesnay, E., 2018. Prediction of activation patterns preceding hallucinations in patients with schizophrenia using machine learning with structured sparsity. Hum Brain Mapp 39, 1777–1788.

Deshpande, G., Hu, X., 2012. Investigating effective brain connectivity from fMRI data: past findings and current issues with reference to Granger causality analysis. Brain Connect 2, 235–245.

Dum, R.P., Strick, P.L., 1991. The origin of corticospinal projections from the premotor areas in the frontal lobe. J. Neurosci. 11, 667–689.

Duval, E.R., Sheynin, J., King, A.P., Phan, K.L., Simon, N.M., Martis, B., Porter, K.E., Norman, S.B., Liberzon, I., Rauch, S.A.M., 2020. Neural function during emotion processing and modulation associated with treatment response in a randomized clinical trial for posttraumatic stress disorder. Depress Anxiety 37, 670–681.

Eickhoff, S.B., Stephan, K.E., Mohlberg, H., Grefkes, C., Fink, G.R., Amunts, K., Zilles, K., 2005. A new SPM toolbox for combining probabilistic cytoarchitectonic maps and functional imaging data. Neuroimage 25, 1325–1335.

Eklund, A., Nichols, T.E., Knutsson, H., 2016. Cluster failure: Why fMRI inferences for spatial extent have inflated false-positive rates. Proc. Natl. Acad. Sci. U.S.A. 113, 7900–7905.

Esteban, O., Markiewicz, C.J., Blair, R.W., Moodie, C.A., Isik, A.I., Erramuzpe, A., Kent, J.D., Goncalves, M., DuPre, E., Snyder, M., Oya, H., Ghosh, S.S., Wright, J., Durnez, J., Poldrack, R.A., Gorgolewski, K.J., 2019. fMRIPrep: a robust preprocessing pipeline for functional MRI. Nat. Methods 16, 111–116.

Etkin, A., Maron-Katz, A., Wu, W., Fonzo, G.A., Huemer, J., Vértes, P.E., Patenaude, B., Richiardi, J., Goodkind, M.S., Keller, C.J., Ramos-Cejudo, J., Zaiko, Y.V., Peng, K.K., Shpigel, E., Longwell, P., Toll, R.T., Thompson, A., Zack, S., Gonzalez, B., Edelstein, R., Chen, J., Akingbade, I., Weiss, E., Hart, R., Mann, S., Durkin, K., Baete, S.H., Boada, F.E., Genfi, A., Autea, J., Newman, J., Oathes, D.J., Lindley, S.E., Abu-Amara, D., Arnow, B.A., Crossley, N., Hallmayer, J., Fossati, S., Rothbaum, B.O., Marmar, C.R., Bullmore, E.T., O’Hara, R., 2019. Using fMRI connectivity to define a treatment-resistant form of post-traumatic stress disorder. Sci Transl Med 11.

Fonzo, G.A., Goodkind, M.S., Oathes, D.J., Zaiko, Y.V., Harvey, M., Peng, K.K., Weiss, M.E., Thompson, A.L., Zack, S.E., Lindley, S.E., Arnow, B.A., Jo, B., Rothbaum, B.O., Etkin, A., 2021. Amygdala and Insula Connectivity Changes Following Psychotherapy for Posttraumatic Stress Disorder: A Randomized Clinical Trial. Biol Psychiatry 89, 857–867.

Fonzo, G.A., Simmons, A.N., Thorp, S.R., Norman, S.B., Paulus, M.P., Stein, M.B., 2010. Exaggerated and disconnected insular-amygdalar blood oxygenation level-dependent response to threat-related emotional faces in women with intimate-partner violence posttraumatic stress disorder. Biol Psychiatry 68, 433–441.

Forman, S.D., Cohen, J.D., Fitzgerald, M., Eddy, W.F., Mintun, M.A., Noll, D.C., 1995. Improved assessment of significant activation in functional magnetic resonance imaging (fMRI): use of a cluster-size threshold. Magn Reson Med 33, 636–647.

Fox, M.D., Snyder, A.Z., Vincent, J.L., Corbetta, M., Van Essen, D.C., Raichle, M.E., 2005. The human brain is intrinsically organized into dynamic, anticorrelated functional networks. Proc. Natl. Acad. Sci. U.S.A. 102, 9673–9678.

Gogolla, N., 2017. The insular cortex. Curr. Biol. 27, R580–R586.

Gorgolewski, K., Burns, C.D., Madison, C., Clark, D., Halchenko, Y.O., Waskom, M.L., Ghosh, S.S., 2011. Nipype: a flexible, lightweight and extensible neuroimaging data processing framework in python. Front Neuroinform 5, 13.

Graziano, M.S.A., Cooke, D.F., 2006. Parieto-frontal interactions, personal space, and defensive behavior. Neuropsychologia 44, 845–859.

Greicius, M.D., Krasnow, B., Reiss, A.L., Menon, V., 2003. Functional connectivity in the resting brain: A network analysis of the default mode hypothesis. Proc Natl Acad Sci U S A 100, 253–258.

Greicius, M.D., Srivastava, G., Reiss, A.L., Menon, V., 2004. Default-mode network activity distinguishes Alzheimer’s disease from healthy aging: evidence from functional MRI. Proc. Natl. Acad. Sci. U.S.A. 101, 4637–4642.

Harricharan, S., Nicholson, A.A., Thome, J., Densmore, M., McKinnon, M.C., Théberge, J., Frewen, P.A., Neufeld, R.W.J., Lanius, R.A., 2020. PTSD and its dissociative subtype through the lens of the insula: Anterior and posterior insula resting-state functional connectivity and its predictive validity using machine learning. Psychophysiology 57, e13472.

Horesh, D., Brown, A.D., 2020. Traumatic stress in the age of COVID-19: A call to close critical gaps and adapt to new realities. Psychol Trauma 12, 331–335.

Kennis, M., Rademaker, A.R., van Rooij, S.J.H., Kahn, R.S., Geuze, E., 2014. Resting state functional connectivity of the anterior cingulate cortex in veterans with and without post-traumatic stress disorder. Hum Brain Mapp 36, 99–109.

Koch, S.B.J., van Zuiden, M., Nawijn, L., Frijling, J.L., Veltman, D.J., Olff, M., 2016. ABERRANT RESTING-STATE BRAIN ACTIVITY IN POSTTRAUMATIC STRESS DISORDER: A META-ANALYSIS AND SYSTEMATIC REVIEW. Depress Anxiety 33, 592–605.

Korgaonkar, M.S., Chakouch, C., Breukelaar, I.A., Erlinger, M., Felmingham, K.L., Forbes, D., Williams, L.M., Bryant, R.A., 2020. Intrinsic connectomes underlying response to trauma-focused psychotherapy in post-traumatic stress disorder. Transl Psychiatry 10, 270.

Kurth, F., Zilles, K., Fox, P.T., Laird, A.R., Eickhoff, S.B., 2010. A link between the systems: functional differentiation and integration within the human insula revealed by meta-analysis. Brain Struct Funct 214, 519–534.

Laird, A.R., Fox, P.M., Eickhoff, S.B., Turner, J.A., Ray, K.L., McKay, D.R., Glahn, D.C., Beckmann, C.F., Smith, S.M., Fox, P.T., 2011. Behavioral interpretations of intrinsic connectivity networks. J Cogn Neurosci 23, 4022–4037.

Larøi, F., Sommer, I.E., Blom, J.D., Fernyhough, C., Ffytche, D.H., Hugdahl, K., Johns, L.C., McCarthy-Jones, S., Preti, A., Raballo, A., Slotema, C.W., Stephane, M., Waters, F., 2012. The characteristic features of auditory verbal hallucinations in clinical and nonclinical groups: state-of-the-art overview and future directions. Schizophr Bull 38, 724–733.

Lefebvre, S., Demeulemeester, M., Leroy, A., Delmaire, C., Lopes, R., Pins, D., Thomas, P., Jardri, R., 2016. Network dynamics during the different stages of hallucinations in schizophrenia. Hum Brain Mapp 37, 2571–2586.

Lewis, S.J., Arseneault, L., Caspi, A., Fisher, H.L., Matthews, T., Moffitt, T.E., Odgers, C.L., Stahl, D., Teng, J.Y., Danese, A., 2019. The epidemiology of trauma and post-traumatic stress disorder in a representative cohort of young people in England and Wales. Lancet Psychiatry 6, 247–256.

Liu, G., Sobering, G., Duyn, J., Moonen, C.T., 1993. A functional MRI technique combining principles of echo-shifting with a train of observations (PRESTO). Magn Reson Med 30, 764–768.

Mahan, A.L., Ressler, K.J., 2012. Fear conditioning, synaptic plasticity and the amygdala: implications for posttraumatic stress disorder. Trends Neurosci 35, 24–35.

Malejko, K., Abler, B., Plener, P.L., Straub, J., 2017. Neural Correlates of Psychotherapeutic Treatment of Post-traumatic Stress Disorder: A Systematic Literature Review. Front Psychiatry 8, 85.

Mary, A., Dayan, J., Leone, G., Postel, C., Fraisse, F., Malle, C., Vallée, T., Klein-Peschanski, C., Viader, F., de la Sayette, V., Peschanski, D., Eustache, F., Gagnepain, P., 2020. Resilience after trauma: The role of memory suppression. Science 367.

Menon, 2015. Salience Network, in: Toga, A.W. (Ed.), Brain Mapping. Academic Press, Waltham, pp. 597–611.

Menon Uddin, L.Q., 2010. Saliency, switching, attention and control: a network model of insula function. Brain Struct Funct 214, 655–667. https://doi.org/10.1007/s00429-010-0262-0

Mesulam, M.-M., Mufson, E.J., 1982. Insula of the old world monkey. Architectonics in the insulo-orbito-temporal component of the paralimbic brain. Journal of Comparative Neurology

Mushtaq, D., Ali, A., Margoob, M.A., Murtaza, I., Andrade, C., 2012. Association between serotonin transporter gene promoter-region polymorphism and 4- and 12-week treatment response to sertraline in posttraumatic stress disorder. J Affect Disord 136, 955–962.

Neggers, S.F.W., Hermans, E.J., Ramsey, N.F., 2008. Enhanced sensitivity with fast three-dimensional blood-oxygen-level-dependent functional MRI: comparison of SENSE-PRESTO and 2D-EPI at 3 T. NMR Biomed 21, 663–676.

Neria, Y., 2021. Functional Neuroimaging in PTSD: From Discovery of Underlying Mechanisms to Addressing Diagnostic Heterogeneity. Am J Psychiatry 178, 128–135.

Newman, M.G., Llera, S.J., Erickson, T.M., Przeworski, A., Castonguay, L.G., 2013. Worry and generalized anxiety disorder: a review and theoretical synthesis of evidence on nature, etiology, mechanisms, and treatment. Annu Rev Clin Psychol 9, 275–297.

Nicholson, A.A., Rabellino, D., Densmore, M., Frewen, P.A., Paret, C., Kluetsch, R., Schmahl, C., Théberge, J., Neufeld, R.W.J., McKinnon, M.C., Reiss, J., Jetly, R., Lanius, R.A., 2016. The neurobiology of emotion regulation in posttraumatic stress disorder: Amygdala downregulation via real-time fMRI neurofeedback. Hum Brain Mapp 38, 541–560.

Nieuwenhuys, R., 2012. Chapter 7 - The insular cortex: A review, in: Hofman, M.A., Falk, D. (Eds.), Progress in Brain Research, Evolution of the Primate Brain. Elsevier, pp. 123–163.

Palaniyappan, L., Liddle, P.F., 2012. Does the salience network play a cardinal role in psychosis? An emerging hypothesis of insular dysfunction. J Psychiatry Neurosci 37, 17–27.

Paus, T., 2001. Primate anterior cingulate cortex: where motor control, drive and cognition interface. Nat. Rev. Neurosci. 2, 417–424.

Rangaprakash, D., Dretsch, M.N., Venkataraman, A., Katz, J.S., Denney, T.S., Deshpande, G., 2018a. Identifying disease foci from static and dynamic effective connectivity networks: Illustration in soldiers with trauma. Hum Brain Mapp 39, 264–287. https://doi.org/10.1002/hbm.23841

Rangaprakash, D., Dretsch, M.N., Venkataraman, A., Katz, J.S., Denney, T.S., Deshpande, G., 2018b. Identifying disease foci from static and dynamic effective connectivity networks: Illustration in soldiers with trauma. Hum Brain Mapp 39, 264–287.

Roebroeck, A., Formisano, E., Goebel, R., 2005. Mapping directed influence over the brain using Granger causality and fMRI. Neuroimage 25, 230–242.

Roullet, P., Vaiva, G., Véry, E., Bourcier, A., Yrondi, A., Dupuch, L., Lamy, P., Thalamas, C., Jasse, L., El Hage, W., Birmes, P., 2021. Traumatic memory reactivation with or without propranolol for PTSD and comorbid MD symptoms: a randomised clinical trial. Neuropsychopharmacology.

Santarnecchi, E., Bossini, L., Vatti, G., Fagiolini, A., La Porta, P., Di Lorenzo, G., Siracusano, A., Rossi, S., Rossi, A., 2019. Psychological and Brain Connectivity Changes Following Trauma-Focused CBT and EMDR Treatment in Single-Episode PTSD Patients. Front Psychol 10, 129.

Sheynin, J., Duval, E.R., King, A.P., Angstadt, M., Phan, K.L., Simon, N.M., Rauch, S.A.M., Liberzon, I., 2020. Associations between resting-state functional connectivity and treatment response in a randomized clinical trial for posttraumatic stress disorder. Depression and Anxiety.

Simmons, A.N., Norman, S.B., Spadoni, A.D., Strigo, I.A., 2013. Neurosubstrates of remission following prolonged exposure therapy in veterans with posttraumatic stress disorder. Psychother Psychosom 82, 382–389.

Sperduti, M., Delaveau, P., Fossati, P., Nadel, J., 2011. Different brain structures related to self- and external-agency attribution: a brief review and meta-analysis. Brain Struct Funct 216, 151–157.

Sridharan, D., Levitin, D.J., Menon, V., 2008. A critical role for the right fronto-insular cortex in switching between central-executive and default-mode networks. PNAS 105, 12569–12574.

Stevens, J.S., Reddy, R., Kim, Y.J., van Rooij, S.J.H., Ely, T.D., Hamann, S., Ressler, K.J., Jovanovic, T., 2018. Episodic memory after trauma exposure: Medial temporal lobe function is positively related to re-experiencing and inversely related to negative affect symptoms. Neuroimage Clin 17, 650–658.

Stoeckel, C., Gough, P.M., Watkins, K.E., Devlin, J.T., 2009. Supramarginal gyrus involvement in visual word recognition. Cortex 45, 1091–1096.

Szeszko, P.R., Yehuda, R., 2019. Magnetic resonance imaging predictors of psychotherapy treatment response in post-traumatic stress disorder: A role for the salience network. Psychiatry Res 277, 52–57.

Thierrée, S., Richa, S., Brunet, A., Egreteau, L., Roig, Q., Clarys, D., El-Hage, W., 2020. Trauma reactivation under propranolol among traumatized Syrian refugee children: preliminary evidence regarding efficacy. Eur J Psychotraumatol 11, 1733248.

Uddin, L.Q., Nomi, J.S., Hebert-Seropian, B., Ghaziri, J., Boucher, O., 2017. Structure and function of the human insula. Journal of clinical neurophysiologylll: official publication of the American Electroencephalographic Society 34, 300.

van den Heuvel, M.P., Sporns, O., 2013. Network hubs in the human brain. Trends Cogn. Sci. (Regul. Ed.) 17, 683–696.

van Gelderen, P., Duyn, J.H., Ramsey, N.F., Liu, G., Moonen, C.T.W., 2012. The PRESTO technique for fMRI. Neuroimage 62, 676–681.

Ventureyra, V.A.G., Yao, S.-N., Cottraux, J., Note, I., De Mey-Guillard, C., 2002. The validation of the Posttraumatic Stress Disorder Checklist Scale in posttraumatic stress disorder and nonclinical subjects. Psychother Psychosom 71, 47–53.

Watkins, E.R., 2008. Constructive and unconstructive repetitive thought. Psychol Bull 134, 163–206.

Weng, Y., Qi, R., Zhang, L., Luo, Y., Ke, J., Xu, Q., Zhong, Y., Li, J., Chen, F., Cao, Z., Lu, G., 2019. Disturbed effective connectivity patterns in an intrinsic triple network model are associated with posttraumatic stress disorder. Neurol. Sci. 40, 339–349.

Yehuda, R., Hoge, C.W., McFarlane, A.C., Vermetten, E., Lanius, R.A., Nievergelt, C.M., Hobfoll, S.E., Koenen, K.C., Neylan, T.C., Hyman, S.E., 2015. Post-traumatic stress disorder. Nat Rev Dis Primers 1, 15057.

Zhu, X., Suarez-Jimenez, B., Lazarov, A., Helpman, L., Papini, S., Lowell, A., Durosky, A., Lindquist, M.A., Markowitz, J.C., Schneier, F., Wager, T.D., Neria, Y., 2018. Exposure-based therapy changes amygdala and hippocampus resting-state functional connectivity in patients with posttraumatic stress disorder. Depress Anxiety 35, 974–984.

