## Supplementary material for "Intrusive Experiences In Post-Traumatic Stress Disorder: Treatment Response Induces Changes In The Effective Connectivity of The Anterior Insula"

***Supplementary table 1.* Distribution of the Index Trauma in the sample**.

|  | | | | |
| --- | --- | --- | --- | --- |
|  | | **Response** | | |
| **Index trauma** | | **no** | | **yes** |
| Combat |  | 1 |  | 0 |
| Life threat |  | 1 |  | 2 |
| Motor vehicle accident, work accident |  | 2 |  | 3 |
| Physical assault |  | 4 |  | 3 |
| Sexual trauma, rape |  | 1 |  | 5 |
| Suicide |  | 2 |  | 0 |
| Terrorist attack |  | 0 |  | 3 |
| Other |  | 3 |  | 0 |

***Supplementary table 2*. Whole-sample *Granger causality analysis* seeded on the anterior insula (AI) at baseline**. Coordinates are reported in the MNI (Montreal Neurological Institute) space.

| **x** | **y** | **z** | **GCM score** | **p-value** | **label** |
| --- | --- | --- | --- | --- | --- |
| 42 | 9 | 3 | 2.343144 | 0.000034 | Central Opercular Cortex |
| 36 | 18 | 27 | 1.625094 | 0.001433 | Middle Frontal Gyrus |
| 27 | -33 | 69 | 1.081782 | 0.008486 | Postcentral Gyrus |
| -3 | -15 | 48 | 1.625094 | 0.000595 | Precentral Gyrus |
| 9 | -48 | 36 | 2.054894 | 0.007480 | Cingulate Gyrus, posterior division |
| -6 | -60 | 54 | 0.781360 | 0.001673 | Precuneus Cortex |
| -10 | -3 | 12 | 1.056062 | 0.010858 | Left Thalamus |
| -30 | 42 | 25 | 1.311266 | 0.011709 | Frontal Pole |
| -30 | 15 | 12 | 2.492637 | 0.000628 | Insular Cortex |

***Supplementary table 3*. Brain regions influenced by the anterior insula (AI) showing a significant difference between V7 and V1 in responders**. Coordinates are reported in the MNI (Montreal Neurological Institute) space.

| **x** | **y** | **z** | **t-value** | **p-value** | **Label** |
| --- | --- | --- | --- | --- | --- |
| 60 | 24 | 21 | -4.157842 | 0.000260 | Inferior Frontal Gyrus, pars triangularis |
| 33 | 12 | -3 | -3.346891 | 0.002274 | Insular cortex |
| 10 | 6 | 57 | -3.259043 | 0.002853 | Juxtapositional Lobule Cortex |
| 3 | 36 | 18 | 2.706012 | 0.011287 | Cingulate Gyrus, anterior division |
| -3 | -9 | 57 | -4.029006 | 0.000370 | Juxtapositional Lobule Cortex |
| -9 | -12 | 48 | -3.627500 | 0.001088 | Juxtapositional Lobule Cortex |
| -48 | -72 | -30 | 2.939507 | 0.006391 | Cerebellum left Crus I |

***Supplementary table 4*. Brain regions influenced by the anterior insula (AI) showing a significant difference between V7 and V1 in non-responders**. Coordinates are reported in the MNI (Montreal Neurological Institute) space.

| **x** | **y** | **z** | **t-value** | **p-value** | **Label** |
| --- | --- | --- | --- | --- | --- |
| 51 | 18 | 42 | -3.656260 | 0.001008 | Middle Frontal Gyrus |
| 51 | -54 | -21 | 2.810747 | 0.008766 | Inferior Temporal Gyrus, temporococipital part |
| 39 | 3 | 12 | 3.456738 | 0.001708 | Central Opercular Cortex |
| 27 | 12 | 3 | 2.947619 | 0.006264 | Right Putamen |
| -12 | -66 | 63 | 3.103135 | 0.004244 | Lateral Occipital Cortex, superior division |
| -12 | -78 | 3 | -3.230455 | 0.003070 | Intracalcarine Cortex |
| -45 | 15 | -3 | 3.454878 | 0.001716 | Frontal Operculum Cortex |
| -54 | -30 | 45 | 3.344527 | 0.002288 | Supramarginal Gyrus, anterior division |
| -64 | -42 | 36 | 2.870195 | 0.007582 | Supramarginal Gyrus, posterior division |

***Supplementary table 5*. Changes in the anterior insula effective connectivity between patients receiving (traumatic memory reactivation + propranolol) and those receiving (traumatic memory reactivation + placebo).** Regions exhibiting a significant difference in *Granger causality analysis* maps between V1 and V7 according to group allocation (propranolol vs placebo). Coordinates are reported in MNI (Montreal Neurological Institute) space.

| **x** | **y** | **z** | **F-value** | **p-value** | **Label** |
| --- | --- | --- | --- | --- | --- |
| 54 | 24 | 12 | 9.314099 | 0.004939 | Inferior Frontal Gyrus |

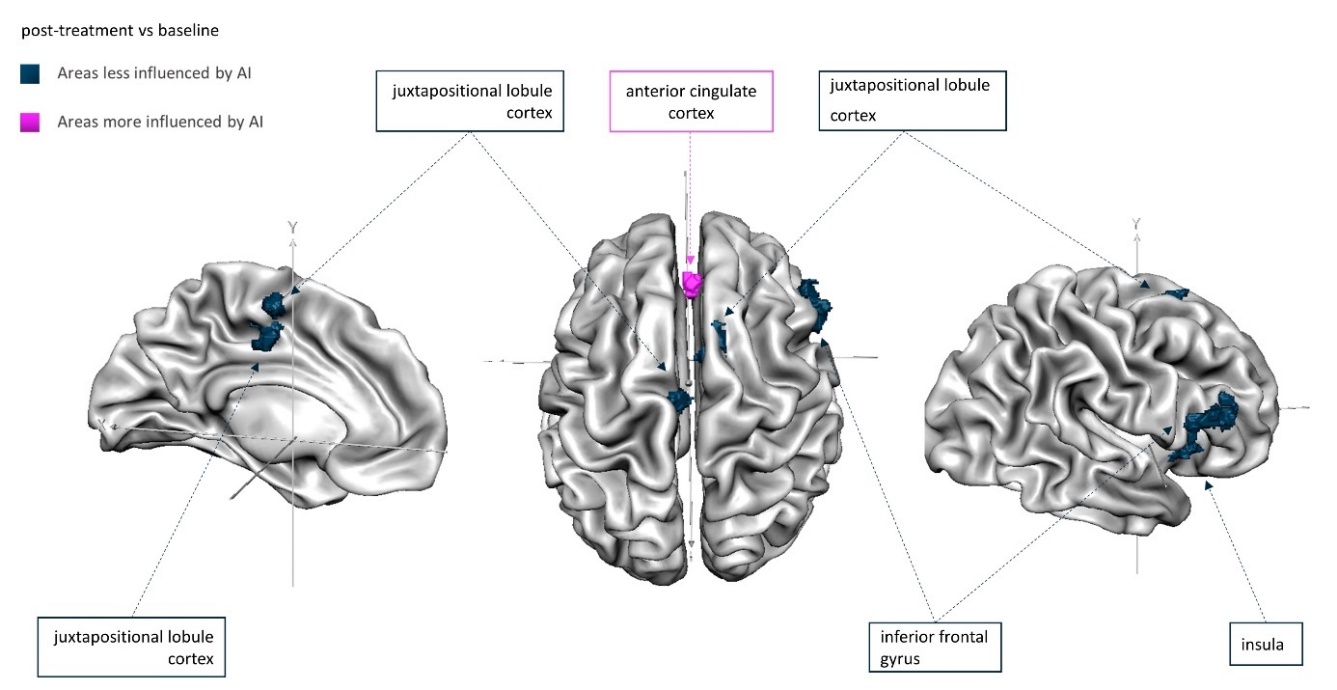
***Supplementary Figure 2.* Changes in *Granger causality* maps seeded on the anterior insula (AI) in responders to therapy (cortical clusters).** Brain areas less influenced by AI after effective treatment are depicted in dark blue. Brain areas more influenced by AI after effective treatment are depicted in pink (i.e. dorsal anterior cingulate cortex, part of the salience network).

***Supplementary Figure 3*. Changes in *Granger Causality* maps seeded on the Anterior Insula (AI) in non-responders to therapy (cortical clusters).** Brain areas less influenced by AI after effective treatment are depicted in dark blue. Brain areas more influenced by AI after effective treatment are depicted in pink.

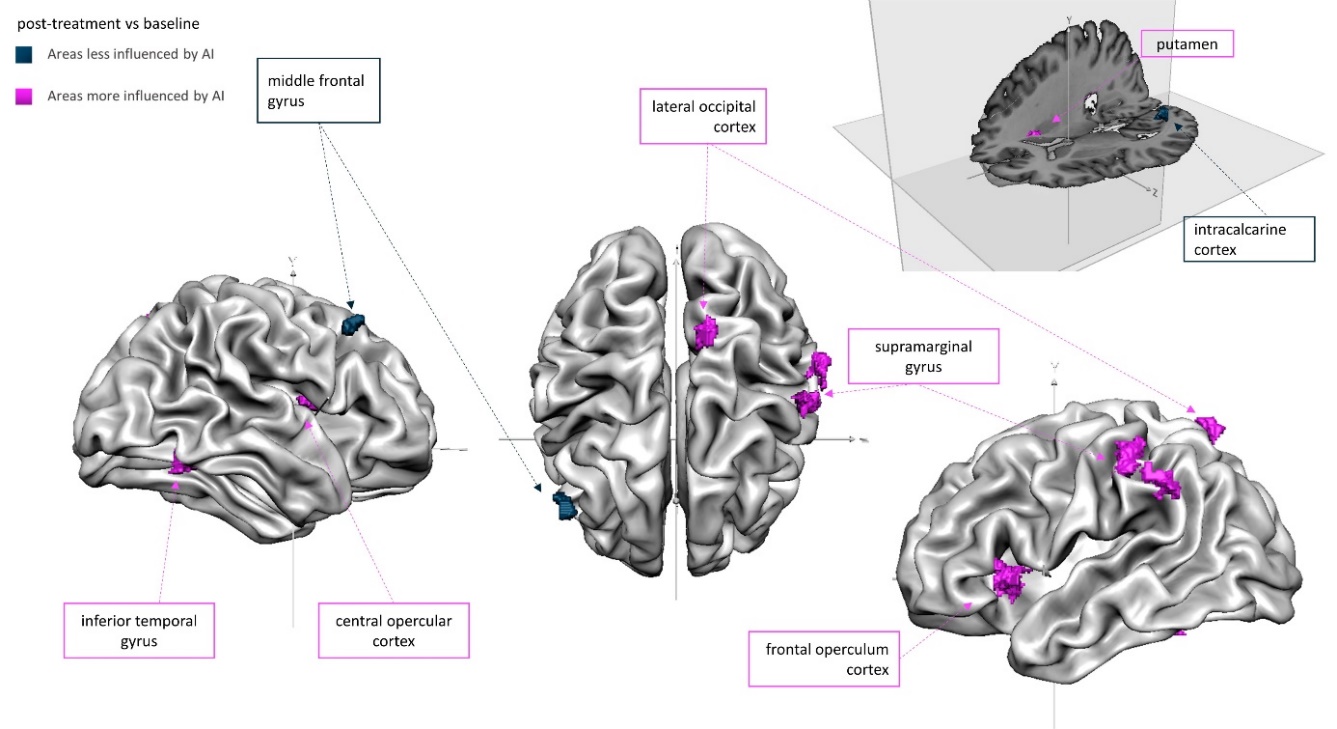

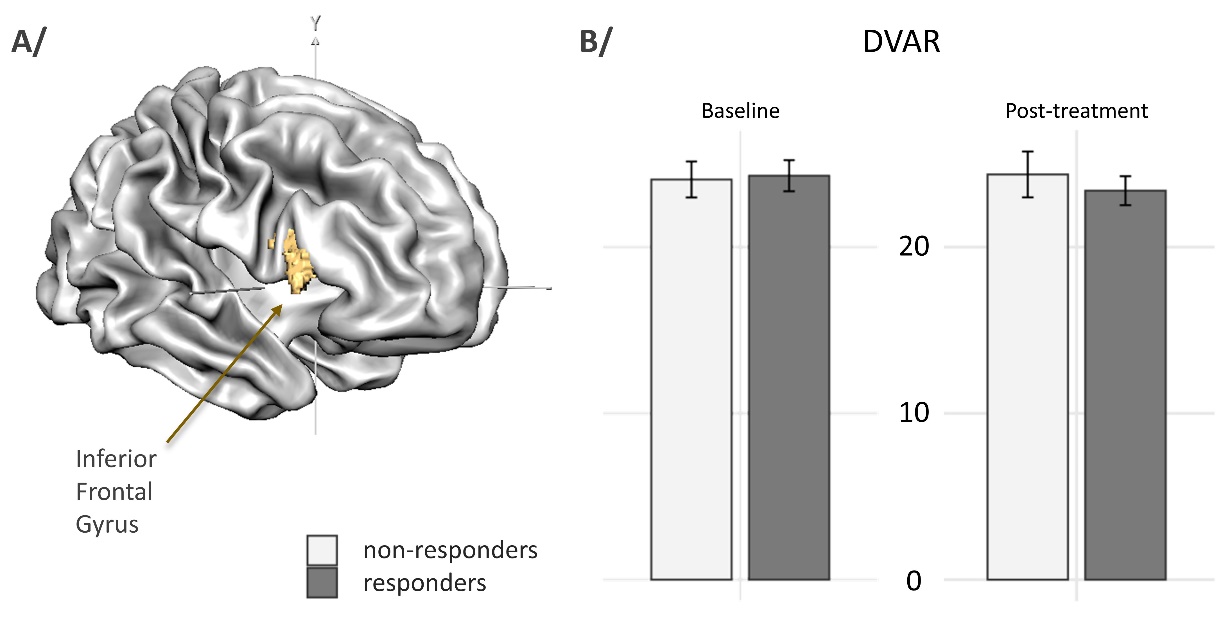
***Supplementary Figure 4*. (A) Brain regions showing a significant (time-point x treatment) interaction for *Granger causality analysis* seeded on the anterior insula (AI)**. The right inferior frontal gyrus was the only area under AI’s influence (shown in yellow), and did not overlap with the network linked with treatment response. **(B)** **Comparison of motion-related fMRI quality metrics**. Responders and non-responders did not differ in terms of fMRI confounds measured pre and post-treatment (DVARs), attesting that head-motion cannot account for the present results.
